# A specific eIF4A paralog facilitates LARP1-mediated translation repression during mTORC1 inhibition

**DOI:** 10.1101/2021.09.18.460932

**Authors:** Yuichi Shichino, Mari Mito, Kazuhiro Kashiwagi, Mari Takahashi, Takuhiro Ito, Nicholas T. Ingolia, Shintaro Iwasaki

## Abstract

Eukaryotic translation initiation factor (eIF) 4A — a DEAD-box RNA-binding protein — plays an essential role in translation initiation. Two mammalian eIF4A paralogs, eIF4A1 and eIF4A2, have been assumed to be redundant because of their high homology, and the difference in their functions has been poorly understood. Here, we show that eIF4A1, but not eIF4A2, enhances translational repression during the inhibition of mechanistic target of rapamycin complex 1 (mTORC1), an essential kinase complex controlling cell proliferation. RNA-immunoprecipitation sequencing (RIP-Seq) of the two eIF4A paralogs revealed that eIF4A1 preferentially binds to mRNAs containing terminal oligopyrimidine (TOP) motifs, whose translation is rapidly repressed upon mTOR inhibition. This biased interaction depends on a La-related RNA-binding protein, LARP1. Ribosome profiling revealed that the deletion of *EIF4A1*, but not *EIF4A2*, rendered the translation of TOP mRNAs resistant to mTOR inactivation. Moreover, eIF4A1 enhances the affinity between TOP mRNAs and LARP1 and thus ensures stronger translation repression upon mTORC1 inhibition. Our data show that the distinct protein interactions of these highly homologous translation factor paralogs shape protein synthesis during mTORC1 inhibition and provide a unique example of the repressive role of a universal translation activator.

## Introduction

Translation initiation — a rate-limiting step of protein synthesis (Shah et al., 2013; Yan et al., 2016) — is precisely controlled to regulate cellular homeostasis and stress responses. Eukaryotic cells harness a number of eukaryotic translation initiation factors (eIFs) to achieve accurate and efficient initiation (Sonenberg and Hinnebusch, 2009). Among the eIFs, the most abundant factor is eIF4A, a DEAD-box RNA-binding protein. eIF4A forms an eIF4F complex together with the 7-methylguanosine (m^7^G) cap-binding factor eIF4E and scaffold protein eIF4G. This complex facilitates the recruitment of the 43S preinitiation complex to the 5′ end of mRNA and the subsequent scanning process along the 5′ untranslated region (UTR) for the recognition of the start codon (Brito Querido et al., 2020; Hinnebusch, 2014). Because eIF4A has ATP-dependent helicase activity (García-García et al., 2015; Rogers et al., 1999; Rozen et al., 1990), it has been thought to function primarily in unwinding secondary structures in the 5′ UTR inhibitory for scanning (Andreou and Klostermeier, 2013; Steinberger et al., 2020; Waldron et al., 2019). However, its pervasive impacts on protein synthesis regardless of the structural complexity of mRNAs indicate a more complex role of eIF4A in translation (Iwasaki et al., 2016; Sen et al., 2015; Sokabe and Fraser, 2017; Yourik et al., 2017).

Mammalian cells possess two highly homologous cytoplasmic eIF4A paralogs, eIF4A1 and eIF4A2, that are 90% identical in amino acid sequence (Nielsen and Trachsel, 1988). Since both paralogs have RNA binding potential (Chen et al., 2021; Chu et al., 2019), are assembled into the eIF4F complex (Fukao et al., 2014; Yoder-Hill et al., 1993), and recruit ribosomes to mRNA (Robert et al., 2020), they have long been assumed to have redundant roles in translation initiation.

However, several lines of evidence have suggested functional differences in eIF4A paralogs. The expression balance between eIF4A1 and eIF4A2 is variable across human tissues; while eIF4A1 is generally dominant, several tissues (*i.e.,* brain and skeletal muscle) express more eIF4A2 than eIF4A1 (Galicia-Vázquez et al., 2012; Nielsen and Trachsel, 1988). Even in the same cell type, cellular status, such as growth arrest or the expression of a specific transcription factor, may alter the equilibrium of the paralogs (Lin et al., 2008; Williams-Hill et al., 1997). More directly, the depletion of eIF4A1, but not eIF4A2, impairs global protein synthesis and subsequent cell growth (Galicia-Vázquez et al., 2012). However, the functional differences between the paralogs remain largely elusive.

The translation program of a cell is highly dependent on nutrient status and dramatically rewired under starvation conditions. Mechanistic target of rapamycin complex 1 (mTORC1), a serine-threonine kinase complex, links nutrient sensing to protein synthesis (Saxton and Sabatini, 2017). The acute inactivation of mTORC1 immediately reduces the rate of translation, particularly of mRNAs with terminal oligopyrimidine (TOP) motifs (defined as a cytosine followed by continuous 4-14 pyrimidines) at the very 5′ end of the mRNA (Hsieh et al., 2012; Jefferies et al., 1994; Levy et al., 1991; Thoreen et al., 2012). Since this motif is found in the mRNAs encoding nearly all ribosome proteins and many translation factors (Meyuhas and Kahan, 2015), the reduced production of translation machinery further extends global translation repression.

The selective repression of TOP mRNA translation upon mTORC1 inhibition is mediated by RNA-binding La-related protein 1 (LARP1) (Fonseca et al., 2015; Lahr et al., 2017; Philippe et al., 2018, 2020; Smith et al., 2020). LARP1 directly binds to the m^7^G cap and pyrimidine sequence via its RNA-binding DM15 domain and competes with eIF4E for m^7^G cap binding (Lahr et al., 2015, 2017; Philippe et al., 2018). In nutrient-rich situations, translational repression by LARP1 is largely suppressed by mTORC1- mediated phosphorylation, which reduces its affinity to the TOP motif (Jia et al., 2021; Philippe et al., 2018). These properties of LARP1 illustrate how TOP mRNAs are selectively repressed during mTORC1 inhibition, although other reports argued that LARP1 acts positively on or is irrelevant to translation (Gentilella et al., 2017; Tcherkezian et al., 2014). The molecular details of how LARP1 represses TOP mRNAs have yet to be fully explored.

In this study, we showed that, counterintuitively, a specific eIF4A paralog functions to repress translation during mTORC1 inhibition by enhancing the interaction of LARP1 with TOP mRNAs. RNA immunoprecipitation-sequencing (RIP-Seq) revealed that the two eIF4A paralogs have different properties in RNA binding and that eIF4A1 has a strong bias toward TOP mRNAs. Using quantitative proteomics, we identified LARP1 as an eIF4A1-selective interactor. LARP1 indeed defines the preferential interaction between eIF4A1 and TOP mRNA. Strikingly, *EIF4A1* knockout (KO) cells, but not *EIF4A2* KO cells, showed resistance in the translation repression of TOP mRNAs and cell growth inhibition upon pharmacological perturbation of mTOR. The interaction between LARP1 and TOP motif is predominantly enhanced by eIF4A1 compared with eIF4A2. Collectively, our data demonstrated that in these highly homologous translation factors, only eIF4A1 differentially modulates the accessibility of LARP1 to TOP mRNA and ensures efficient translation repression in mTORC1-inhibited conditions. This study provides an example in which a positive regulator of translation has an alternative mode to inhibit translation.

## Results

### eIF4A1 preferentially interacts with TOP mRNAs

To characterize the mRNA-binding preference of the two eIF4A paralogs, we set out RNA immunoprecipitation-sequencing (RIP-Seq) from RNA complexed with eIF4A1 and eIF4A2 in human embryonic kidney (HEK) 293T cells (Figure 1A). The disproportionate expression of endogenous eIF4A1 and eIF4A2 (with eIF4A1 expressed at much higher levels) (Figure S1A) may hamper the comparison of RIP-seq data from these two proteins. Thus, we generated stable cell lines expressing streptavidin-binding peptide (SBP)-tagged eIF4A1 or eIF4A2 driven by a tetracycline-inducible promoter. These cell lines showed comparable expression of eIF4A paralogs (Figure S1B), enabling us to compare their RNA preferences quantitatively. We purified SBP-tagged eIF4A paralogs (Figure S1C) and associated mRNAs for downstream RNA-Seq analyses.

**Figure 1.**
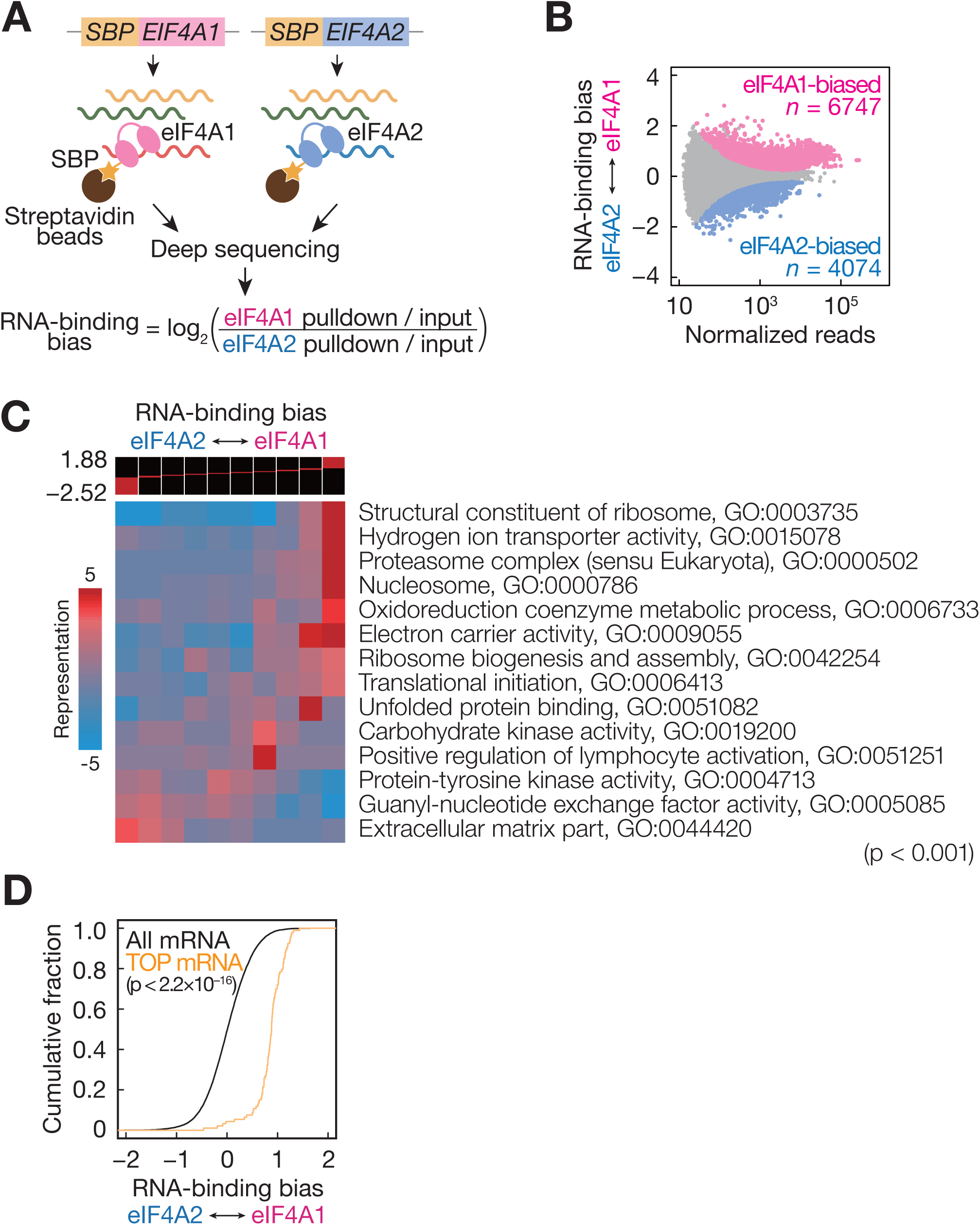
eIF4A1 preferentially interacts with TOP mRNAs. (A) Schematic of RIP-Seq analysis and calculation for RNA-binding bias between eIF4A paralogs. (B) MA (M, log ratio; A, mean average) plot of the RNA-binding bias. eIF4A1-biased and eIF4A2-biased RNAs (false discovery rate [FDR] < 0.05) were highlighted. (C) GO analysis of RNA-binding bias visualized by iPAGE (Goodarzi et al., 2009). (D) Cumulative distribution of the RNA-binding bias for TOP mRNAs compared to all mRNAs. The p-value was calculated by Mann-Whitney *U* test. See also Figure S1 and Table S1.

To estimate the RNA preferences of the eIF4A paralogs, we calculated the over- or under-representation of RNAs in the complexes over input RNAs (*i.e.*, relative eIF4A occupancy) (Figure 1A). We observed that both eIF4A paralogs have diverse biases across the transcriptome (Figure S1D). Whereas the correspondence of mRNA preference between eIF4A paralogs was observed (Figure S1D), a subset of mRNAs bound to eIF4A paralogs in a biased manner. The “RNA-binding bias”, calculated as a ratio of occupancy of eIF4A1 to eIF4A2 (Figure 1A), was used to quantitatively assess the difference in binding preferences and revealed thousands of transcripts with biased affinity toward either paralog (Figure 1B and Table S1). Therefore, eIF4A1 and eIF4A2 have differential mRNA binding characteristics.

To investigate the functional implication of this RNA-binding bias, we performed Gene Ontology (GO) analysis and found enriched terms in both eIF4A1- and eIF4A2-biased mRNAs (Figure 1C). In particular, mRNAs encoding ribosome proteins preferentially bound to eIF4A1 over eIF4A2. Notably, these mRNAs have a 5′ terminal oligopyrimidine (TOP) motif, which consists of a C directly adjacent to the 5′ cap and subsequent pyrimidine nucleotides (Meyuhas and Kahan, 2015). Indeed, the classical TOP mRNAs (defined in (Meyuhas and Kahan, 2015)) were predominantly associated with eIF4A1 (Figure 1D).

### eIF4A1 preferentially interacts with LARP1

Given that DEAD-box RNA binding proteins in general binds to the ribose-phosphate backbone of RNA without base recognition (Linder and Jankowsky, 2011), we hypothesized that the biased binding to TOP mRNAs might be conferred by another protein that preferentially binds to eIF4A1. To quantitatively and comprehensively explore proteins associated with eIF4A paralogs, we conducted mass spectrometry with stable isotope labeling by amino acids in cell culture (SILAC) (Emmott and Goodfellow, 2014), followed by pulldown of each eIF4A paralog from HEK293T cells (Figure 2A). After continuous culture of cells in media containing different sets of isotopes, we purified each SBP-tagged paralog, pooled the isolated proteins, and measured the ratio of isotope incorporation using liquid chromatography-tandem mass spectrometry (LC-MS/MS). A majority of proteins, including translation initiation factors, interacted with both paralogs equivalently (Figure 2B, S2A, and Table S2). We noted that eIF4G1 showed a modest bias to eIF4A1, but eIF4G2 did not (Figure S2A), consistent with an earlier report (Sugiyama et al., 2017). Similarly, we found that 14 and 7 proteins had stronger affinities for eIF4A1 and eIF4A2, respectively (Figure 2B).

**Figure 2.**
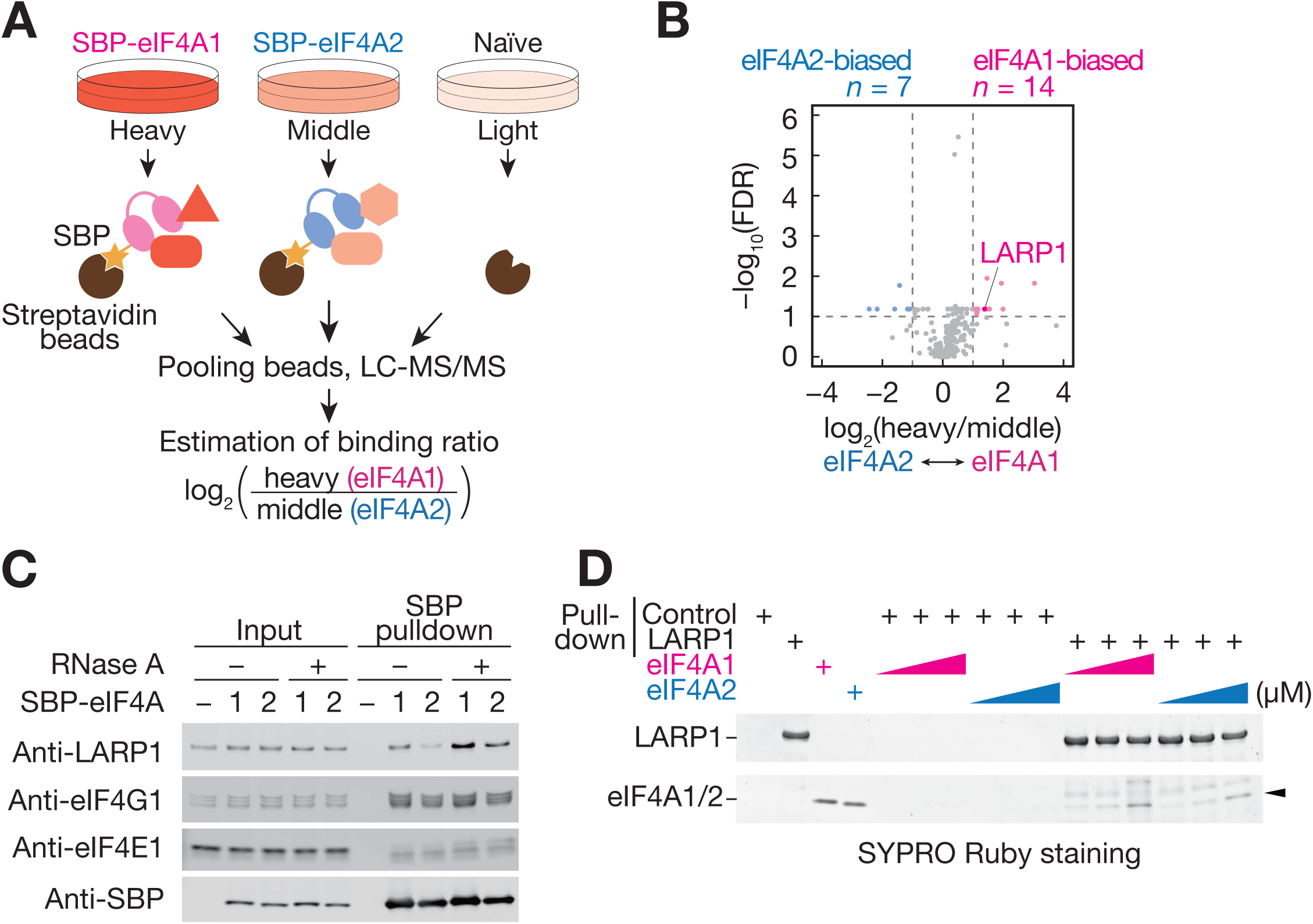
LARP1 shows preferential binding to eIF4A1. (A) Schematic of SILAC-based mass spectrometry analysis and estimation of the binding ratio between eIF4A paralogs. (B) Volcano plot of protein-binding bias. eIF4A1- and eIF4A2-biased proteins (defined as FDR < 0.1 and log2[heavy/middle] > 1 and < -1, respectively) are highlighted. (C) Western blotting for proteins associated with SBP-tagged eIF4A proteins. (D) *In vitro* pulldown assay between purified LARP1 and recombinant eIF4A paralogs. The beads with or without FLAG-tagged LARP1 were incubated with 0.2, 0.6, and 2.0 µM of His-tagged eIF4A1 or eIF4A2. For loading control, 0.2 µM of eIF4A proteins were used. Proteins were stained using SYPRO Ruby. Arrowhead indicates eIF4A1/2 protein. See also Figure S2 and Table S2.

Among the eIF4A1-biased proteins, we found La-related protein 1 (LARP1), which recognizes the 5′ cap structure and the subsequent TOP motif (Figure 2B) (Lahr et al., 2017; Philippe et al., 2018). Western blotting confirmed the biased interaction of LARP1 with eIF4A1 over eIF4A2 (Figure 2C). The preferential interaction of LARP1 with eIF4A1 was maintained even after RNase treatment (Figure 2C and S2B), indicating that the LARP1-eIF4A1 complex is formed through protein-protein interactions. The direct interaction between LARP1 and eIF4A1 was further confirmed by pulldown assay with recombinant proteins (Figure 2D). Importantly, even in this purified system, LARP1 still showed stronger affinity to eIF4A1 than eIF4A2 (Figure 2D).

Then, we investigated the regions responsible for the interaction of eIF4A1 with LARP1. Although the amino acid sequences of the eIF4A paralogs are highly similar overall, the N-terminal 12-13 amino acids are relatively variable (designated as the N-terminal variable region, or NVR) (Figure S2C). However, swapping the NVRs of the paralogs did not alter the preference of LARP1 for eIF4A1 (Figure S2D). We then focused on the helicase core, consisting of N-terminal and C-terminal domains. Isolated N- or C-terminal domains from eIF4A1 (and eIF4A2) did not interact with LARP1 (Figure S2E), suggesting that both domains of eIF4A1 are necessary for LARP1 binding.

### LARP1 facilitates the biased interaction of eIF4A1 with TOP mRNAs

Given the interaction of eIF4A1 with LARP1 (Figure 2) and of LARP1 with TOP motif RNA (Fonseca et al., 2015; Lahr et al., 2017; Philippe et al., 2018, 2020), we speculated that LARP1 determines the biased interaction between eIF4A1 and TOP mRNAs. To test this model, we performed RIP-Seq from SBP-tagged eIF4A paralogs upon *LARP1* knockdown by small interfering RNA (siRNA) (Figure S3A and S3B). The depletion of *LARP1* significantly reduced the interaction of eIF4A1 with TOP mRNAs (Figure 3A), whereas the interactions with other mRNAs without TOP motifs were only modestly affected (Figure 3B). Thus, LARP1 ensures the association of eIF4A1 with TOP mRNAs.

**Figure 3.**
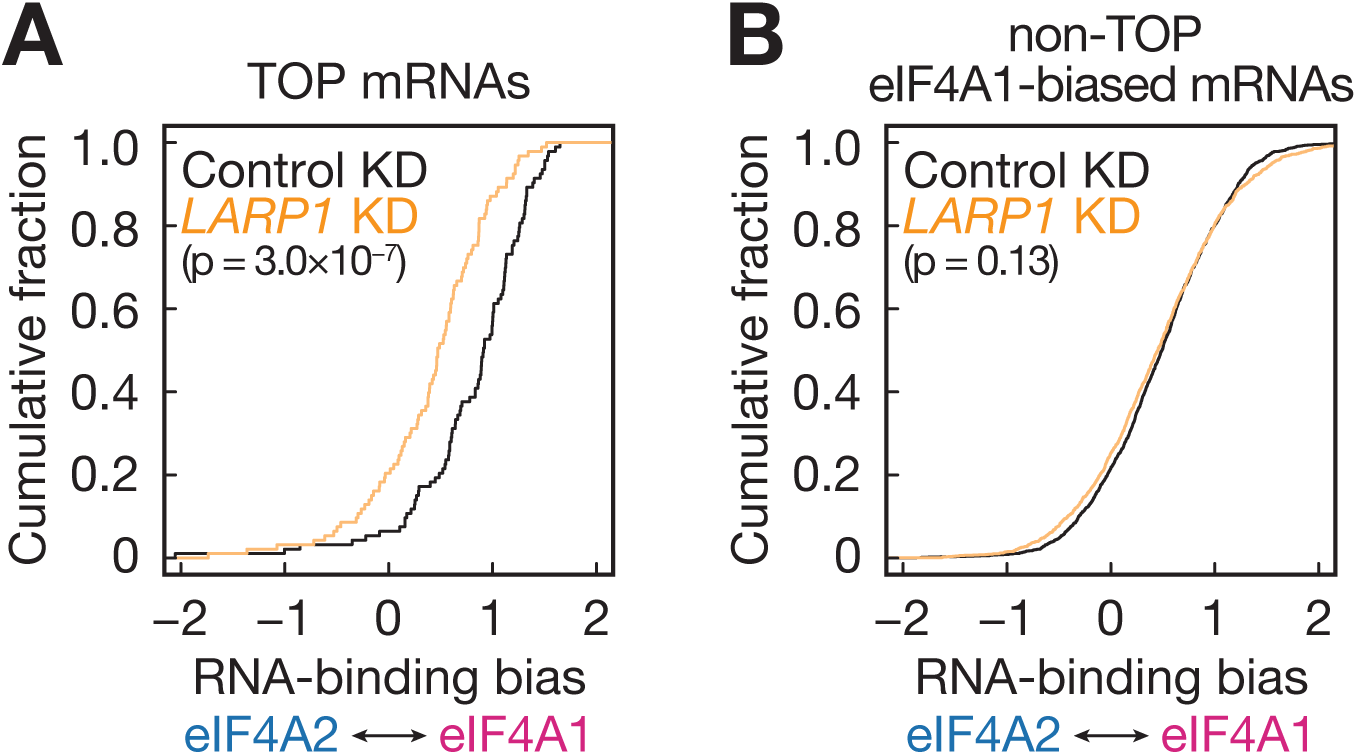
Depletion of LARP1 eliminates the biased interaction between eIF4A1 and TOP mRNAs. (A) Cumulative distribution of the RNA-binding bias (Figure 1A) of TOP mRNAs in *LARP1* or control knockdown cells. (B) Same as (A) but for eIF4A1-biased mRNAs without TOP motifs. The p-values were calculated by Mann-Whitney *U* test. See also Figure S3.

### Two eIF4A paralogs differentially shape translation profiles

To investigate how eIF4A paralogs impact translation, we generated *EIF4A1* and *EIF4A2* KO cell lines. For the knockout of *EIF4A1*, we used the SBP-eIF4A1 [Phe163Leu- Ile199Met] *eIF4A1^em1SINI^* cell line (Iwasaki et al., 2019), in which all endogenous *EIF4A1* alleles have been disrupted by CRISPR/Cas9 and the exogenous *SBP-EIF4A1* under tetracycline-inducible promoter complimented the loss of the endogenous *EIF4A1*. The continuous culture of this cell line in tetracycline-free media for 10 days or more gradually suppressed the expression of exogenous *SBP-EIF4A1* and ultimately led to the depletion of all the eIF4A1 in the cells (Figure S4A) (denoted this cell as *EIF4A1* KO). We also applied the CRISPR/Cas9 technique to the *EIF4A2* gene and introduced mutations that cause premature termination codons in both alleles (Figure S4B) (denoted *EIF4A2* KO). Western blotting using specific antibodies against the N-terminal, isoform-specific regions confirmed the null expression of the paralogs in these cell lines (Figure 4A). Notably, the expression of eIF4A2 was upregulated in *EIF4A1* KO cells, consistent with earlier observations of *EIF4A1* knockdown by siRNA (Galicia-Vázquez et al., 2012). Both KO cell lines showed growth retardation (Figure S4C) and a reduction in global protein synthesis (Figure 4B), as measured by metabolic labeling of newly synthesized proteins with *O*-propargyl-puromycin (OP-puro) (Liu et al., 2012). In both assays, we observed more severe effects in *EIF4A1* KO cells than in *EIF4A2* KO cells. These results indicate that both paralogs have pivotal roles in translation in growing HEK293T cells.

**Figure 4.**
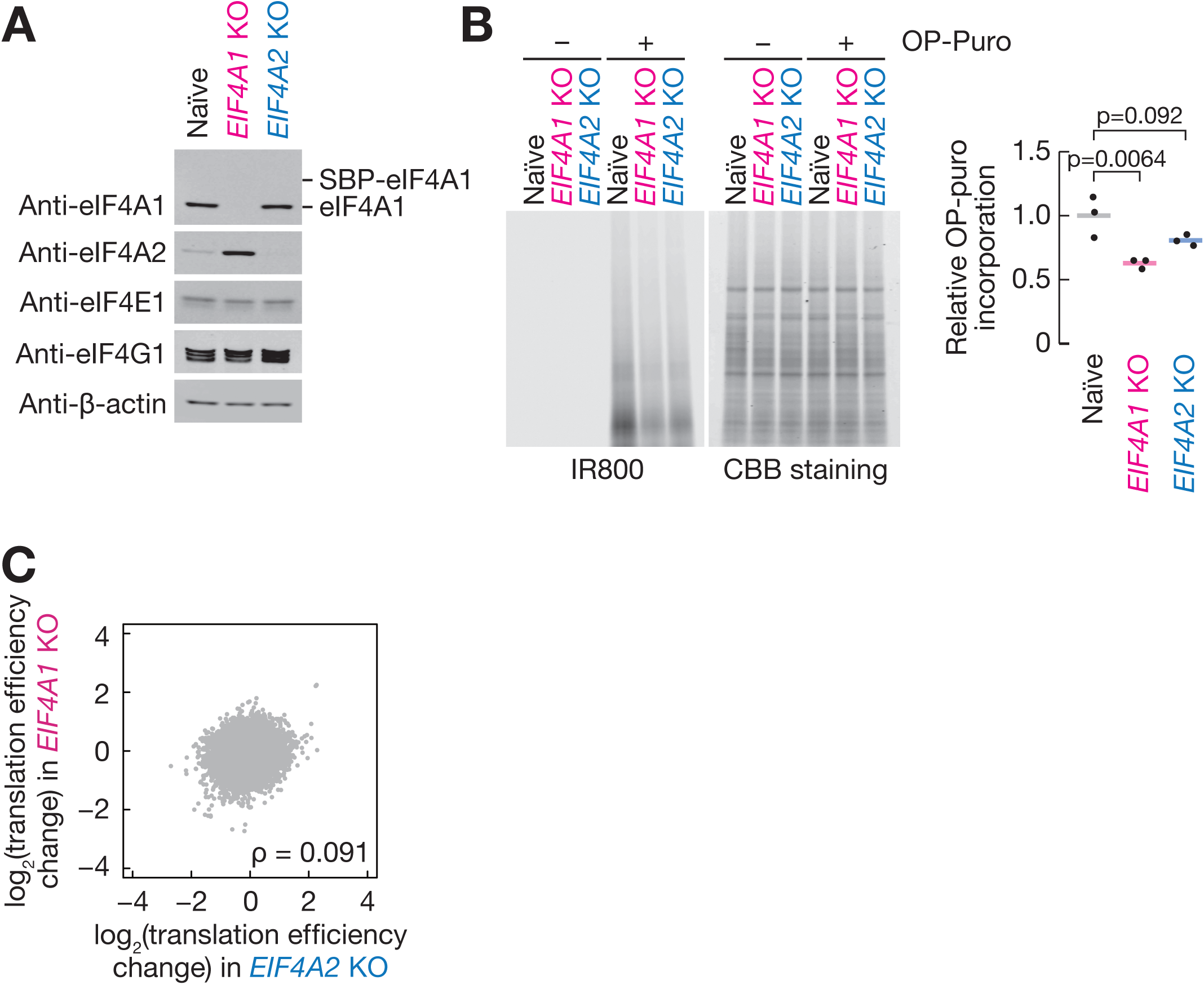
eIF4A1 and eIF4A2 target different mRNAs in translation. (A) Western blotting of eIF4A1, eIF4A2, eIF4E1, and eIF4G1 in naïve, *EIF4A1* KO, and *EIF4A2* KO cells. β-Actin was used as the loading control. (B) Global protein synthesis rate in *EIF4A1* KO and *EIF4A2* KO cells as measured by OP-puro labeling. Quantification of nascent proteins (infrared 800 signal) normalized to total protein (signal of CBB staining) is shown on the right. Data from three replicates (points) and the mean (bar) are shown. The p-values were calculated by Dunnett’s multiple comparison test. (C) Comparison of translation efficiency changes in *EIF4A1* KO and *EIF4A2* KO cells. ρ, Spearman’s rank correlation coefficient. See also Figure S4 and Table S3.

Ribosome profiling of both KO cell lines revealed divergent alterations in translation status. We found that the translation efficiency varied across the transcriptome upon deletion of both eIF4A paralogs (Figure S4D, S4E, and Table S3). However, the trends were poorly correlated (Figure 4C), suggesting that the two eIF4A paralogs drive protein synthesis from different mRNAs in cells.

### eIF4A1 is required for high sensitivity to mTOR inhibition

The translation of TOP mRNAs is repressed selectively and intensively upon mTORC1 inhibition (Hsieh et al., 2012; Thoreen et al., 2012) through their interaction with LARP1 (Fonseca et al., 2015; Lahr et al., 2017; Philippe et al., 2018). Considering that LARP1 and eIF4A1 form a complex, we hypothesized that eIF4A1 may play a role in LARP1- mediated translation repression during mTORC1 inhibition. To test this idea, we performed ribosome profiling after treatment with PP242, a catalytic mTOR inhibitor (Table S4). As reported previously (Iwasaki et al., 2016), PP242 hampered protein synthesis from TOP mRNAs (Figure S5A). However, the efficacy of this inhibition was attenuated in *EIF4A1* KO cells, irrespective of the concentration of PP242 (Figure 5A and S5B). In contrast, *EIF4A2* depletion did not impact TOP mRNA translation at low PP242 doses (Figure S5B) and even further repressed their translation at high doses (Figure 5A). This phenotype in *EIF4A1* KO cells could not be explained by the modulation of LARP1 expression or by insensitivity of mTOR to PP242 (Figure S5C), which was monitored by the dephosphorylation of eIF4E-binding protein 1 (4EBP1), a direct target of mTORC1 (Saxton and Sabatini, 2017).

**Figure 5.**
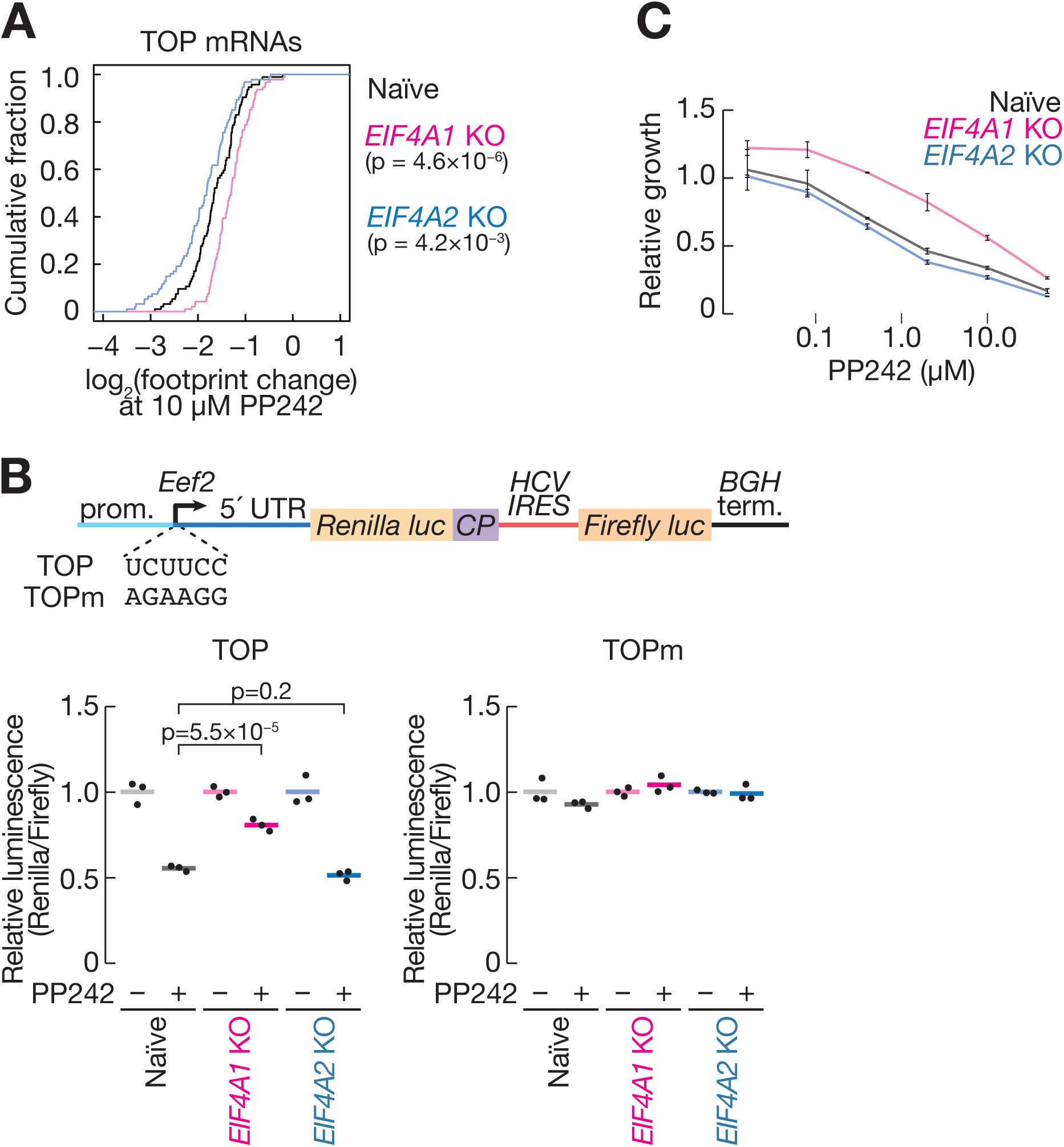
*EIF4A1* KO cells are resistant to mTOR inhibition. (A) Cumulative distribution of TOP mRNAs along translation changes upon treatment with 10 µM PP242 in naïve, *EIF4A1* KO, and *EIF4A2* KO cells. The p-values were calculated by Mann-Whitney *U* test. (B) (Top) Schematic representation of reporter genes. TOP-containing *Eef2* 5′ UTR and its endogenous promoter were conjugated with *Renilla* luciferase with the destabilizing sequence (CP). Firefly luciferase with HCV IRES was used as an internal control. In the TOPm reporter, the 6 nucleotides surrounding the transcription start site of *Eef2* (UCUUCC) were substituted by AGAAGG. (Bottom) Relative luminescence of TOP and TOPm reporters in naïve, *EIF4A1* KO, and *EIF4A2* KO cells treated with 3 µM PP242 for 2 h. The *Renilla* luciferase signal was normalized to that of firefly luciferase. Data from three replicates (points) and the mean (bars) are shown. The p-values were calculated by Dunnett’s multiple comparison test. (C) Relative growth rate of naïve, *EIF4A1* KO, and *EIF4A2* KO cells treated with PP242 for 22 h. Signals were normalized to those in control DMSO-treated samples. See also Figure S5 and Table S4.

Although we performed RIP-Seq and mass spectrometry from eIF4A paralogs in growing cells (Figure 1 and 2), the protein-RNA complex was largely maintained even during mTOR inhibition. We confirmed that the preferential complex formation of LARP1-eIF4A1 still occurred upon PP242 treatment (Figure S5D). Similarly, even with mTOR inhibition, endogenous TOP mRNAs, *e.g., RPL13* and *RPS27A*, showed similar RNA-binding bias to eIF4A1 (Figure S5E and S5F). Thus, the biochemical properties of the eIF4A1-LARP1-TOP mRNA complex are maintained even during mTORC1 inactivation.

We then tested the importance of TOP motifs for eIF4A1-facilitated translation repression with a reporter assay. Here we harness a polycistronic translation system (Figure 5B, top), in which 5′ *Renilla* luciferase is translated by TOP motif 5′ UTR and 3′ firefly luciferase by an eIF4A-independent internal ribosome entry site (IRES) from Hepatitis C virus (HCV) (Hellen and Sarnow, 2001) for the normalization. Consistent with the ribosome profiling results, the repression of TOP mRNA reporters by PP242 was dampened in the absence of eIF4A1 (Figure 5B, bottom). In contrast, the mutated TOP (TOPm) reporter was insensitive to mTOR inhibition in all three cell lines.

Since TOP motifs are found in mRNAs encoding ribosome proteins, the suppression of protein synthesis from those mRNAs directly limits cell growth (Saxton and Sabatini, 2017). mTOR inhibition by PP242 reduced cell growth; however, *EIF4A1* deletion attenuated the rate (Figure 5C), whereas *EIF4A2* KO cells showed PP242 sensitivity comparable to that of naïve cells. Thus, we concluded that eIF4A1, but not eIF4A2, is required for the full translational repression of TOP mRNAs and thereby for cell growth control upon mTOR inhibition.

### eIF4A1 enhances the interaction between LARP1 and TOP mRNAs

How does eIF4A1 lead to stronger translation repression by LARP1? As LARP1 directly binds to the cap structure and following TOP motif to block translation (Lahr et al., 2017; Philippe et al., 2018), we speculated that eIF4A1 might enhance the affinity of LARP1 to the TOP motif to ensure strong repression. Here, we monitored the association of LARP1 (and eIF4A1/2 proteins) with TOP motif RNA by ultraviolet (UV) crosslinking. Purified LARP1 and eIF4A paralogs were incubated with radiolabeled short RNAs bearing the cap and the TOP motif and then crosslinked to RNA with UV; sodium dodecyl sulfate polyacrylamide gel electrophoresis (SDS-PAGE) was then used to resolve these complexes. By this method, we recapitulated the specific binding of LARP1 to the TOP motif, as pyrimidines but not purines adjacent to the m^7^G cap structure allowed strong crosslinking to the LARP1 protein (Figure S6A) (Lahr et al., 2017; Philippe et al., 2018). Moreover, dephosphorylated LARP1 protein purified from PP242-treated cells showed enhanced binding to TOP RNA (Figure S6A and S6B), as reported previously (Jia et al., 2021; Philippe et al., 2018). We therefore used dephosphorylated LARP1 for further experiments. Supplementation with recombinant eIF4A1 led to a stronger interaction between LARP1 and TOP RNA in a dose-dependent manner (Figure 6A and 6B). eIF4A2 also enhanced the interaction between LARP1 and TOP RNA but to a lesser extent than eIF4A1.

**Figure 6.**
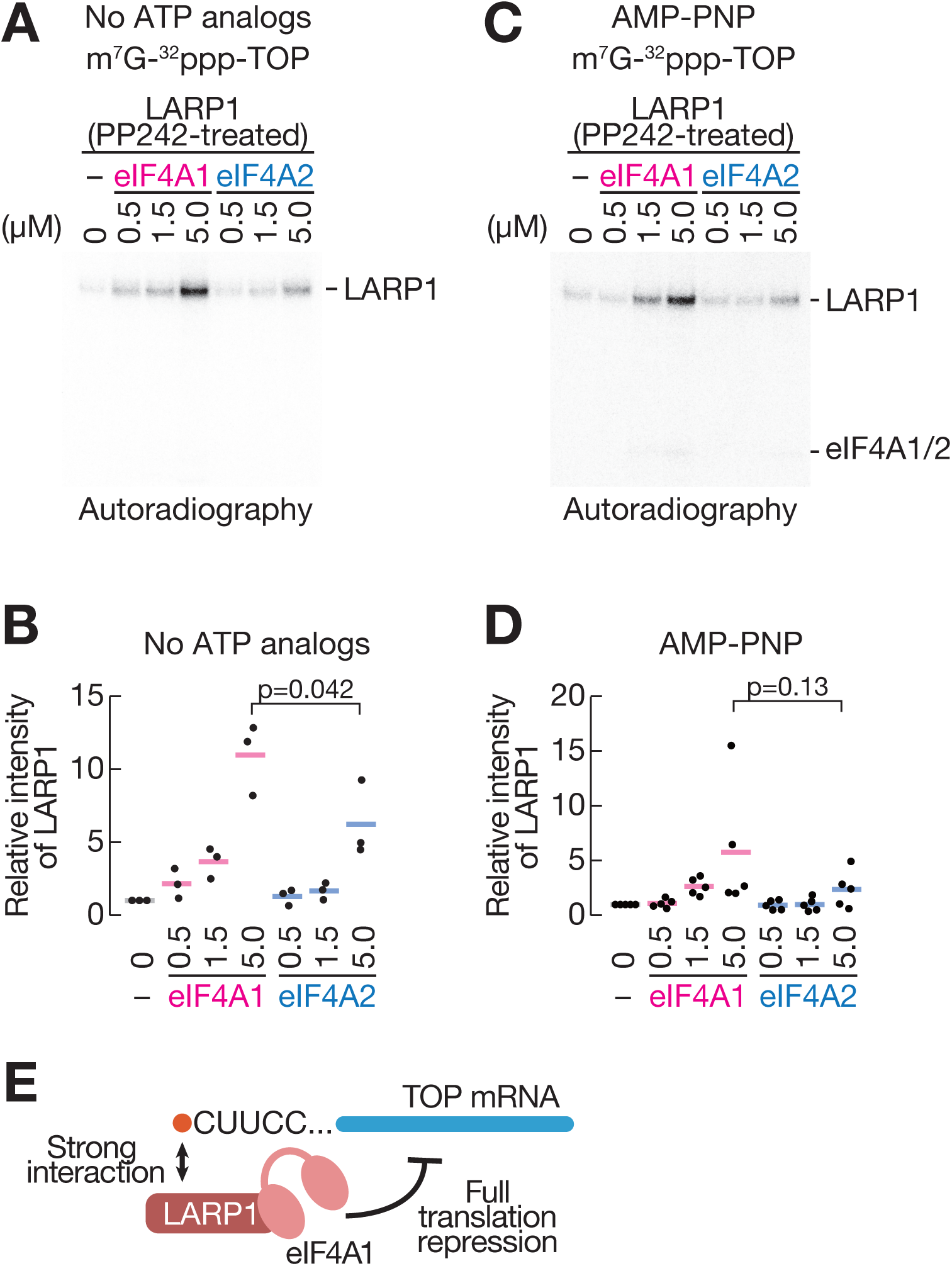
eIF4A1 enhances the interaction between LARP1 and TOP RNA. (A) RNA crosslinking assay between recombinant LARP1 protein and cap-labeled TOP RNA in the presence of eIF4A paralog proteins. Each reaction contained 0.4 µM cap-labeled TOP RNA, 0.25 µM 3×FLAG-6×His-LARP1 purified from PP242-treated cells, and 6×His-eIF4A paralogs at the indicated concentration. (B) Quantification of LARP1 band intensity in (A). Data from three replicates (points) and the mean (bar) are shown. The p-values were calculated by Student’s *t* test. (C) Same as (A) but in the presence of AMP-PNP. (D) Quantification of LARP1 band intensity in (C). (E) Schematic representation of eIF4A1-specific facilitation of translation repression from TOP mRNAs. See also Figure S6.

We noted that, although eIF4A1 is also an RNA binding protein, its RNA association is not a prerequisite for the enhanced LARP1-TOP RNA interaction. Our crosslinking assay did not contain ATP, which is required for eIF4A to bind RNA (Linder and Jankowsky, 2011) (Figure 6A and 6B). Also, we observed essentially the same increase of LARP1 affinity to TOP RNA, even in the presence of AMP-PNP, a ground state ATP analog that fixes eIF4A in RNA binding conformation (Figure 6C and 6D).

Taken together, all the data in our study demonstrated that eIF4A1, but not eIF4A2, has a repressive function, serving to increase the affinity of LARP1 to the TOP motif for full translation repression (Figure 6E).

## Discussion

Since eIF4A has been recognized as a universal driver of protein synthesis, its repressive role in translation, as observed in this study, was unexpected. However, given that compounds, such as rocaglates, may convert eIF4A into a translational repressor (Iwasaki et al., 2016, 2019), the capacity for dual effects on translation regulation may be intrinsic in eIF4A. Through the investigation of the difference between eIF4A1 and eIF4A2, we provided an example of a repressive mode of eIF4A1 in the absence of pharmacological treatment.

Such a binary role of the translation factor has been exemplified in the α subunit of eIF2 heterotrimer. eIF2 complex possessing unphosphorylated eIF2α delivers initiator methionyl-tRNA to the 40S ribosome subunits in a GTP-dependent manner to drive translation initiation (Merrick and Pavitt, 2018). In contrast, stress-induced phosphorylation of eIF2α converts the protein into translation repressor (Wek, 2018), especially as an inhibitor of eIF2B (Kashiwagi et al., 2019; Kenner et al., 2019), a heterodecamer for guanine nucleotide exchange factor for eIF2, leading to global reduction of protein synthesis (so called integrated stress response, ISR). Irrespective of the mRNA selectivity (global vs. TOP mRNA-selective), the stress-evoked repressive mechanisms of general translation factors eIF2α and eIF4A1 may represent a common means to cope with the environmental alterations.

Our study demonstrated that the function of eIF4A1 in protein synthesis may depend on the interaction partners. It is widely recognized that eIF4A1 complexed with eIF4G and eIF4E (*i.e.*, eIF4F complex) should drive general translation, even in mTORC1-inhibited condition. Simultaneously, the large excess of eIF4A1 relative to eIF4G and eIF4E (Figure S1A) allows a subpopulation of eIF4A1 that is free of eIF4G-eIF4E to associate with LARP1 and exert the repressive function. Such an eIF4F-free function of eIF4A1 was also demonstrated for the modulation of RNA contents in the stress granule (Tauber et al., 2020), exemplifying the multi-faceted character of eIF4A1.

Earlier works have implicated eIF4A2 in miRNA-mediated translation repression (Meijer et al., 2013; Wilczynska et al., 2019), although this model has been contested (Fukao et al., 2014; Galicia-Vázquez et al., 2015). In our ribosome profiling data, we did not find any evidence of a role of eIF4A2 in miRNA-mediated translation repression; the translation efficiency of mRNAs targeted by miR-10, the most abundant miRNA in HEK293 cells (Panwar et al., 2017), was not increased but was rather reduced in *EIF4A2* KO cells (Figure S4F). Thus, in addition to the individual reporters (Fukao et al., 2014; Galicia-Vázquez et al., 2015), transcriptome-wide impacts of eIF4A2 in miRNA-mediated translational repression was quite limited.

How eIF4A1 enhances the affinity of LARP1 to TOP mRNAs remains an open question. Molecular dynamics simulations suggested that the binding pocket in the DM15 domain of LARP1 is highly flexible, such that the accessibility of the m^7^G cap and the first C nucleotide in TOP mRNA could be dynamically tunable (Cassidy et al., 2019). Thus, eIF4A1 binding may change the conformation of the DM15 domain to a form with high affinity for the TOP motif.

mTOR inhibitors are promising antitumor drugs (Hua et al., 2019). Since cancer cells show differential sensitivity to mTOR inhibitors (Ducker et al., 2014; Jastrzebski et al., 2018), the prediction of treatment efficacy is a demanding task. Indeed, recent studies have revealed various factors associated with mTOR inhibitor resistance, such as the level of mTOR kinase activity (Rodrik-Outmezguine et al., 2016), the amount of 4EBP1 (Ducker et al., 2014; Jastrzebski et al., 2018), and the expression balance of eIF4E/4EBP1 (Alain et al., 2012). Our data demonstrated that the expression ratio of two eIF4A paralogs could be an additional determinant of mTOR inhibitors to impact in cancer therapy.

## Acknowledgments

We are grateful to all the members of the Iwasaki laboratory for constructive discussions, technical help, and critical reading of the manuscript. We also thank Kosuke Dodo and Mikiko Sodeoka for the imaging scanner Pharos FX, the Support Unit for Bio-Material Analysis, RIKEN CBS Research Resources Division for mass spectrometry and Sanger sequencing and the HOKUSAI SailingShip supercomputer facility at RIKEN for computation support. S.I. was supported by the Ministry of Education, Culture, Sports, Science and Technology (MEXT) (a Grant-in-Aid for Transformative Research Areas [B] “Parametric Translation”, JP20H05784), the Japan Society for the Promotion of Science (JSPS) (a Grant-in-Aid for Young Scientists [A], JP17H04998; a Challenging Research [Exploratory], JP19K22406), AMED (AMED-CREST, JP21gm1410001), and RIKEN (Pioneering Projects “Biology of Intracellular Environments” and Aging Project). T.I. was supported by JSPS (a Grants-in-Aid for Scientific Research (B), JP19H03172), AMED (AMED-CREST, JP21gm1410001), and RIKEN (the BDR Structural Cell Biology Project, the Pioneering Projects “Dynamic Structural Biology” and “Biology of Intracellular Environments”, and the All RIKEN Research Project “Integrated life science research to challenge super aging society”). K.K. was supported by JSPS (a Grant-in-Aid for Early-Career Scientists, JP18K14644). Y.S. was supported by JSPS (a Grant-in-Aid for JSPS Fellows, JP19J00920; a Grant-in-Aid for Early-Career Scientists, JP21K15023) and RIKEN (Special Postdoctoral Researchers and Incentive Research Projects). DNA libraries were sequenced by the Vincent J. Coates Genomics Sequencing Laboratory at UC Berkeley, which is supported by the National Institutes for Health (NIH) Instrumentation Grant (S10 OD018174). Y.S. was a recipient of a JSPS Research Fellow (PD) and the RIKEN Special Postdoctoral Researchers Program.

## Author contributions

Conceptualization, Y.S. and S.I.; Methodology, Y.S. and S.I.; Formal analysis, Y.S.; Investigation, Y.S., M.M., K.K., M.T., and S.I.; Resources, N.T.I.; Writing – Original Draft, Y.S. and S.I.; Writing – Review & Editing, Y.S., M.M., K.K., M.T., T.I., N.T.I., and S.I.; Visualization, Y.S.; Supervision, T.I., N.T.I., and S.I.; Funding Acquisition, K.K., T.I., Y.S., and S.I.

## Experimental procedures

### Cell lines

HEK293 Flp-In T-Rex cells (Thermo Fisher Scientific, R78007) were cultured in DMEM + GlutaMAX-I (Thermo Fisher Scientific, 10566016) with 10% fetal bovine serum (FBS) (Sigma-Aldrich, F7524) at 5% CO2 and 37°C. Tetracycline-free FBS (Biowest, S182T-500) was used for the SBP-eIF4A1 [Phe163Leu-Ile199Met] *eIF4A1^em1SINI^* cell line (Iwasaki et al., 2016) to repress the exogenous expression of SBP-eIF4A1.

Stable integrants of pcDNA5/FRT/TO plasmids were established by cotransfection with pOG44 by X-tremeGENE9 (Roche, 06365787001) and selected with blasticidin S (InvivoGen, ant-bl-1) and hygromycin B (InvivoGen, ant-hg-1). To induce the expression of the integrated genes, cells were cultured in medium containing 1 µg/ml tetracycline for 3 days. The stable cell line possessing SBP-eIF4A1 was previously reported (Iwasaki et al., 2016).

For knockdown of *LARP1*, cells were transfected with siRNAs targeting *LARP1* (ON-TARGETplus Human LARP1 [23367] siRNA SMARTpool, L-027187-00-0020) or control siRNAs (siGENOME Non-Targeting siRNA Pool #1, D-001206-13-20) using the *Trans*IT-X2 Dynamic Delivery System (Mirus Bio, MIR6003) for 72 h.

The *EIF4A2* KO cell line was generated by the CRISPR/Cas9 approach (Iwasaki et al., 2019). Two guide RNAs were transcribed from the template PCR fragments (5′- TAATACGACTCACTATAGG<u>GTGGCTCCGCGGATTATAAC</u>GTTTAAGAGCTAT GCTGGAAACAGCATAGCAAGTTTAAATAAGGCTAGTCCGTTATCAACTTGAAAAAGTGGCACCGAGTCGGTGCTTTTTTT-3′ and 5′- TAATACGACTCACTATAGG<u>CCAGAGGGAATGGACCCCGA</u>GTTTAAGAGCTA TGCTGGAAACAGCATAGCAAGTTTAAATAAGGCTAGTCCGTTATCAACTTG AAAAAGTGGCACCGAGTCGGTGCTTTTTTT-3′, where the underlined characters represent sequences complementary to the *EIF4A2* gene locus) by a T7-Scribe Standard RNA IVT kit (Cellscript, C-AS3107). The guide RNAs and recombinant Cas9-NLS protein (Thermo Fisher Scientific, B25642) were transfected into HEK293 T-Rex cells with Lipofectamine CRISPRMAX Cas9 Transfection Reagent (Thermo Fisher Scientific, CMAX00008). After 48 h of incubation, cells were seeded at 0.1 cells/well in 96-well plates to isolate clonal populations. Cells were screened by sequencing around the targeted genome region and then further confirmed the null expression of eIF4A2 by Western blotting.

PP242 (Sigma-Aldrich, P0037) was added to the medium 30 min before cell lysis unless otherwise noted.

### DNA construction

All PCR amplification and insertion into plasmids were performed with PrimeSTAR MAX and In-Fusion HD (TaKaRa), respectively.

### pcDNA5/FRT/TO-SBP-eIF4A2

A DNA fragment encoding the full-length *EIF4A2* gene (NM_001967) was PCR-amplified from the HEK293T genomic DNA and inserted into pcDNA5/FRT/TO-SBP- eIF4A1 (Iwasaki et al., 2016), replacing the eIF4A1 gene with the eIF4A2 gene.

### pcDNA5/FRT/TO-SBP-eIF4A1-2NVR and SBP-eIF4A2-1NVR

To construct pcDNA5/FRT/TO-SBP-eIF4A1-2NVR, a DNA fragment containing NVR of *EIF4A2* was PCR-amplified from pcDNA5/FRT/TO-SBP-eIF4A2 and inserted into pcDNA5/FRT/TO-SBP-eIF4A1, replacing the NVR of eIF4A1 with the NVR of eIF4A2. pcDNA5/FRT/TO-SBP-eIF4A2-1NVR was also prepared in the same manner.

### pcDNA5/FRT/TO-SBP-eIF4A1-Nterm, SBP-eIF4A2-Nterm, SBP-eIF4A1-Cterm, and SBP-eIF4A2-Cterm

To construct pcDNA5/FRT/TO-SBP-eIF4A1-Nterm and SBP-eIF4A2-Nterm, the N-terminal regions of eIF4A1 and eIF4A2 were PCR-amplified from pcDNA5/FRT/TO- SBP-eIF4A1 and eIF4A2 and inserted into the pcDNA5/FRT/TO-SBP backbone. For pcDNA5/FRT/TO-SBP-eIF4A1-Cterm and SBP-eIF4A2-Cterm, pcDNA5/FRT/TO- SBP-eIF4A1 and eIF4A2 were linearized by PCR, skipping NVRs and N-terminal regions, and self-ligated.

### pcDNA5/FRT/TO-3×FLAG-6×His-LARP1

A DNA fragment encoding the full-length *LARP1* gene (NM_033551) was PCR-amplified from HEK293T genomic DNA and inserted into the pColdI vector (TaKaRa, 3361) at the NdeI and HindIII sites. Using this plasmid as a template, the 6×His-LARP1 sequence was amplified and inserted into pcDNA5/FRT/TO together with a 3×FLAG DNA fragment to construct pcDNA5/FRT/TO-3×FLAG-6×His-LARP1.

### psiCHECK2-Eef2TOP-RLCP-HCVIRES-FL and Eef2TOPm-RLCP-HCVIRES-FL

The TOP and TOPm reporter plasmids were designed as previously described (Philippe et al., 2018), with modifications. A DNA fragment spanning the promoter and the 5′ UTR of *Eef2* was PCR-amplified from mouse NIH3T3 genomic DNA and substituted for the CMV promoter of the psiCHECK2 vector (Promega, C802A). The intergenic region between *Renilla* and firefly luciferase in this plasmid was replaced by the sequence of HCV IRES PCR-amplified from psiCHECK2-HCV-IRES (Iwasaki et al., 2016). A DNA fragment of hCL/hPEST (CP) sequence (Rubio et al., 2014) was inserted downstream of *Renilla* luciferase. The TOPm substitution was introduced by site-directed mutagenesis.

### RIP-Seq

RIP-Seq was performed as described previously (Iwasaki et al., 2016) with modifications. Briefly, cells expressing SBP-eIF4A1 and SBP-eIF4A2 were cultured in a 10-cm dish and lysed with 600 µl of lysis buffer (20 mM Tris-HCl [pH 7.5], 150 mM NaCl, 5 mM MgCl2, 1 mM dithiothreitol [DTT], and 1% Triton X-100). After TURBO DNase treatment (Thermo Fisher Scientific, AM2238) and clarification by centrifugation, the lysate was incubated with 60 µl of lysis buffer-equilibrated Dynabeads M-270 Streptavidin (Thermo Fisher Scientific, 65305) for 30 min at 4°C. The beads were washed five times with lysis buffer containing 1 M NaCl and incubated with 25 µl of biotin elution buffer (lysis buffer with 5 mM D-biotin [Thermo Fisher Scientific, B20656], omitting Triton X-100) for 30 min at 4°C. Half of the eluate was subjected to SDS-PAGE and Coomassie Brilliant Blue (CBB) staining by EzStain AQua (ATTO, AE-1340). The remaining eluate was used for RNA extraction by TRIzol LS (Thermo Fisher Scientific, 10296-010) and a Direct-zol RNA Microprep Kit (Zymo Research, R2060). The RNA-Seq library was prepared by the Ribo-Zero Gold rRNA Removal Kit (Human/Mouse/Rat) (Illumina, RZG1224) and TruSeq Stranded mRNA Library Prep Kit (Illumina, 20020594). The libraries were sequenced on a HiSeq 4000 (Illumina).

For RIP-Seq of *LARP1-*knockdown cells, the lysate was incubated with Dynabeads M-270 Streptavidin for 1 h at 4°C. The beads were washed five times with lysis buffer containing 500 mM NaCl and incubated with 100 µl of biotin elution buffer for 3 h at 4°C.

### Ribosome profiling and RNA-Seq

Ribosome profiling was performed according to a previously described protocol (McGlincy and Ingolia, 2017; Mito et al., 2020). Ribosome-protected RNA fragments ranging from 26-34 nt were gel-excised.

For RNA-Seq, 1 µg of RNA extracted from the same lysate sample with TRIzol LS and a Direct-zol RNA Microprep Kit was used for library preparation as described in the RIP-Seq section. Libraries were sequenced on a HiSeq 4000 instrument (Illumina).

### Deep sequencing data analysis

Data were processed as previously described (McGlincy and Ingolia, 2017) with modifications. Read quality filtering and adapter trimming were performed with Fastp (Chen et al., 2018). After removing noncoding RNA-mapped reads, the remaining reads were aligned to the human genome hg38 and assigned to the GENCODE Human release 32 reference using STAR 2.7.0a (Dobin et al., 2013). For ribosome profiling, the offsets of the A site from the 5′ end of ribosome footprints were determined to be 15 for 26-29 nt, 16 for 30-31 nt, and 17 for 32 nt. For RNA-Seq and RIP-Seq analysis, an offset of 15 was used for all mRNA fragments.

For the calculation of RNA-binding bias, counts from pulldown samples were normalized to those of input samples with the DESeq2 package (Love et al., 2014). The significance was calculated by a likelihood ratio test in a generalized linear model. Translation efficiency, which is an over- or under-representation of ribosome profiling counts over RNA-seq counts, was calculated in a manner similar to that used for RNA-binding bias. Reads corresponding to the first and last five codons in the CDS were excluded from the calculation. Gene ontology analysis was performed with iPAGE (Goodarzi et al., 2009). Target prediction of miRNA was performed with TargetScan 7.2 (Agarwal et al., 2015).

All custom scripts used in this study are available upon request.

### SILAC, SBP pulldown, and mass spectrometry

SILAC was performed with a SILAC Protein Quantitation Kit (LysC) and DMEM (Thermo Fisher Scientific, A33969). Cell lines with SBP-eIF4A1/2 integration were grown in media containing different isotopes (L-lysine-^13^C6,^15^N2 and L-arginine- ^13^C6,^15^N4 for SBP-eIF4A1, L-lysine-4,4,5,5-D4 and L-arginine-^13^C6 for SBP-eIF4A2). Naïve cells were grown in media containing L-lysine and L-arginine. After culture for 2 weeks in these media, cells were cultured for 3 days with 1 µg/ml tetracycline in a 10-cm dish and lysed with 600 µl of lysis buffer. Equal amounts of lysate were incubated with 60 µl of lysis buffer-equilibrated Dynabeads M-280 Streptavidin (Thermo Fisher Scientific, 11206D) for 2 h at 4°C. The beads were washed five times with 150 µl of lysis buffer without Triton X-100. Samples labeled with different isotopes were pooled.

The proteins on beads were reduced by DTT, alkylated with iodoacetamide, and digested with modified trypsin (Wiśniewski et al., 2009). The resulting peptides were subjected to LC-MS/MS using EASY-nLC 1000 (Thermo Fisher Scientific, LC120) and Q Exactive (Thermo Fisher Scientific, IQLAAEGAAPFALGMAZR) equipped with a nanospray ion source. The peptides were separated with a NANO-HPLC C18 capillary column (0.075 x 150 mm, 3 µm, Nikkyo Technos) using a 60 min gradient at a flow rate of 300 nL/min: 5-35% B in 48 min, and then 35-65% B in 12 min (solvent A; 0.1% formic acid, solvent B; 100% CH3CN/0.1% formic acid). The MS and MS/MS data were searched against the Swiss-Prot database using Proteome Discoverer 2.4 (Thermo Fisher Scientific) with MASCOT search engine software (Matrix Science). Proteins with peptide counts greater than 1 were used for analysis. Additionally, proteins with (heavy/light) < 1 or (middle/light) < 1 were excluded because they were pulled down nonspecifically.

### SBP pulldown assay for Western blotting

Cells expressing SBP-tagged eIF4A paralogs and their variants were cultured in a 10-cm dish and lysed with 600 µl of lysis buffer. The lysate (300 µl) was incubated with 30 µl of lysis buffer-equilibrated Dynabeads M-280 Streptavidin for 1 h at 4°C. The beads were washed three times with lysis buffer and subjected to Western blotting. In Figure 2C, lysates were incubated with 10 µg of RNase A (Nacalai, 30100-31) for 1 h at 25°C prior to purification. RNA was extracted with TRIzol LS and a Direct-zol RNA Microprep Kit and examined for digestion by a MultiNA microchip electrophoresis system (Shimadzu, MCE-202).

### Western blotting

Anti-eIF4A1 (Cell Signaling Technology [CST], 12782), anti-eIF4A2 (Abcam, ab104375), anti-eIF4G (CST, 2498), anti-eIF4E (CST, 9742), anti-LARP1 D8J4F (CST, 70180), anti-SBP (Santa Cruz, SB19-C4, sc-101595), anti-4EBP1 (CST, 9452), anti-phosphorylated (Ser65) 4EBP1 (CST, 174A9, 9456), and anti-β-actin (Medical & Biological Laboratories [MBL], M177-3) were used as primary antibodies. IRDye 800CW anti-rabbit IgG (LI-COR, 926-32211) and IRDye 680RD anti-mouse IgG (LI-COR, 925-68070) were used as secondary antibodies. Images were acquired by ODYSSEY CLx (LI-COR).

### RT-qPCR

RNAs bound to SBP-tagged eIF4A paralogs were purified as described above in *RIP-Seq* for *LARP1* knockdown. The RNA was assayed with ReverTra Ace qPCR Master Mix (TOYOBO, FSQ-201) in a Thermal Cycler Dice (TaKaRa) with TB Green Premix Ex Taq II (Tli RNaseH Plus) (TaKaRa, RR820). The primers used in this study were as follows: *RPL13* forward (5′-CAGCGGCTGAAGGAGTACC-3′), *RPL13* reverse (5′-GGTGGCCAGTTTCAGTTCTT-3′), *RPS27A* forward (5′-CCTGATCAGCAGAGACTGATCTT-3′), and *RPS27A* reverse (5′-TTTTCTTAGCACCACCACGA-3′).

### Reporter assay

psiCHECK2-Eef2TOP-RLCP-HCVIRES-FL and psiCHECK2-Eef2TOPm-RLCP-HCVIRES-FL plasmids were transfected into cells with *Trans*IT-293 Reagent (Mirus, MIR2704). After 24 h of incubation, cells were treated with 3 µM PP242 for 2 h and lysed with Passive Lysis Buffer (Promega, E194A). Luminescence was detected with a Dual-Luciferase Reporter Assay System (Promega, E1960) in GloMax Navigator (Promega, GM2000).

### Purification of recombinant eIF4A1 and eIF4A2 proteins

Recombinant eIF4A1 and eIF4A2 proteins were purified as described previously (Chen et al., 2021) with modifications. *E. coli* One Shot BL21 Star (DE3) chemically competent cells (Thermo Fisher Scientific, 44-0049) transformed with pColdI-eIF4A1 or eIF4A2 (Chen et al., 2021; Iwasaki et al., 2019) were cultivated to OD600 0.5 at 37°C in 1 l LB medium with 50 µg/ml ampicillin and then incubated at 15°C overnight with 1 mM IPTG. Cells were lysed by sonication in His lysis buffer (20 mM HEPES-NaOH pH 7.5, 500 mM NaCl, 10 mM imidazole, 10 mM β-mercaptoethanol [ME], and 0.5% NP-40) and clarified by centrifugation. The supernatant containing His-eIF4A1 or His-eIF4A2 was incubated with a 1.5 ml bed volume of Ni-NTA Superflow (Qiagen, 30430) or a 0.75 ml bed volume of Ni-NTA agarose (Qiagen, 30210) for 1 h, respectively. The beads loaded on a gravity column were washed twice with 25 ml of high-salt wash buffer (20 mM HEPES-NaOH pH 7.5, 1 M NaCl, 20 mM imidazole, and 10 mM β-ME) and then twice with 25 ml of the low-salt wash buffer (contents as in the high-salt buffer but with 10 mM NaCl). The His-tagged proteins were finally eluted in 8 ml of elution buffer (20 mM HEPES-NaOH pH 7.5, 10 mM NaCl, 250 mM imidazole, 10 mM β-ME, and 10% glycerol) overnight. Using the NGC Chromatography System (Bio-Rad), the protein was loaded onto a 1 ml HiTrap Heparin HP column (GE Healthcare, 17-0406-01) and eluted with a linear gradient from buffer A (20 mM HEPES-NaOH pH 7.5, 10 mM NaCl, 1 mM DTT, and 10% glycerol) to buffer B (contents as in buffer A but with 1 M NaCl). The peaked fractions were collected, buffer-exchanged in protein storage buffer (20 mM HEPES-NaOH pH 7.5, 100 mM KCl, 1 mM DTT, and 10% glycerol) on a PD-10 column (GE Healthcare, 17085101), concentrated with a 10 kDa MWCO Amicon Ultra-4 device (Millipore, UFC8010), flash-frozen in liquid nitrogen, and stored at -80°C. All purification steps were performed at 4°C.

### Purification of recombinant LARP1 proteins

LARP1 protein was purified as described previously (Philippe et al., 2018) with modifications. Briefly, HEK293 Flp-In T-Rex cells with 3×FLAG-6×His-LARP1 integration were seeded in five 15-cm dishes and incubated with tetracycline for 3 days to induce protein expression. Cells on each dish were washed once with 10 ml of ice-cold PBS and lysed with 1 ml of lysis buffer. After clarification by centrifugation, lysates were incubated with 200 μl of anti-FLAG M2 affinity gel (Sigma-Aldrich, A2220) (pre-equilibrated with lysis buffer) for 1 h at 4°C. The beads were washed five times with 10 ml of FLAG wash buffer 1 (lysis buffer with 1 M NaCl), washed five times with 10 ml of FLAG wash buffer 2 (FLAG wash buffer 1 without Triton X-100), and then incubated with 150 μl of FLAG elution buffer (FLAG wash buffer 2 with 1 mg/ml 3×FLAG peptide [Protein Ark, GEN-3XFLAG-25]) overnight. Subsequently, FLAG-tagged LARP1 was eluted again in two rounds of incubation with 150 µl of FLAG elution buffer for 1 h. Eluted proteins were exchanged into protein storage buffer on an NAP-5 column (GE Healthcare, 17085301), concentrated with a 50 kDa MWCO Amicon Ultra-0.5 device (Millipore, UFC505096), flash-frozen in liquid nitrogen, and stored at -80°C. All purification steps were performed at 4°C.

### In vitro pulldown assay

Cells expressing 3×FLAG-6×His-LARP1 were cultured in a 10-cm dish and lysed with 600 µl of lysis buffer. After clarification, 300 µl of lysates were incubated with 30 μl of Anti-FLAG M2 Magnetic Beads (Sigma-Aldrich, M8823) equilibrated with a lysis buffer for 1 h at 4°C. For the negative control, lysates from naïve cells were used. The beads were washed five times with 500 µl of FLAG wash buffer 1, five times with 500 µl of FLAG wash buffer 2, and three times with a binding buffer [20 mM HEPES-NaOH (pH 7.5), 100 mM KCl, 1 mM DTT, and 1% Triton X-100]. In each washing step, the buffer-suspended beads were incubated for 5 min at 4°C with rotation. After washing, the beads were incubated with the recombinant His-eIF4A1 or eIF4A2 in 10 µl of the binding buffer for 30 min at 37°C, washed three times with 150 µl of the binding buffer, and incubated with 15 µl of the FLAG elution buffer for overnight at 4°C. The eluates were subjected to SDS-PAGE. The gel image was obtained by SYPRO Ruby protein gel stain (Thermo Fisher Scientific, S12001) and Pharos FX (Bio-rad).

### RNA crosslink assay

Triphosphorylated RNA oligonucleotides (TOP, 5′-pppCUCUUCCGCCGU-3′; TOPm, 5′-pppGAGAAGGGCCGU-3′) were synthesized by Nippon Gene. These RNAs were cap-labeled with α-^32^P-GTP (Perkin Elmer, NEG006H), ScriptCap m^7^G Capping System (Cellscript, C-SCCE0625), and ScriptCap 2′-O-Methyltransferase (Cellscript, C- SCMT0625) and purified by a Direct-zol RNA Microprep Kit (Zymo Research, D2060). The radiolabeled TOP RNAs (4 pmol) were incubated with LARP1 protein and eIF4A paralog in 10 µl of the protein storage buffer for 30 min at 37°C. The TOPm RNA was diluted to the same signal intensity as TOP RNA. For the assays with ATP analogs, 1 mM MgCl2 and 1 mM AMP-PNP were included in the reactions. After crosslinking by 30 mJ/cm^2^ of 254 nm UV, proteins were separated by SDS-PAGE. Autoradiographic image was obtained using an Amersham Typhoon Scanner IP system (GE Healthcare).

### Nascent peptide labeling by OP-puro

OP-puro labeling was performed as described previously (Iwasaki et al., 2019). The gel images were acquired and quantified by an ODYSSEY CLx instrument.

### Cell viability assay

Cells were plated on a white 96-well plate (Grenier, 655083) and incubated with RealTime-Glo MT Cell Viability Assay reagent (Promega, G9712). The luminescence was detected by GloMax Navigator.

### Accession numbers

The RIP-Seq, ribosome profiling, and RNA-Seq (GSE126298 and GSE184247) data used in this study were deposited in the National Center for Biotechnology Information (NCBI).

## Figure and table legends

**Figure S1.**
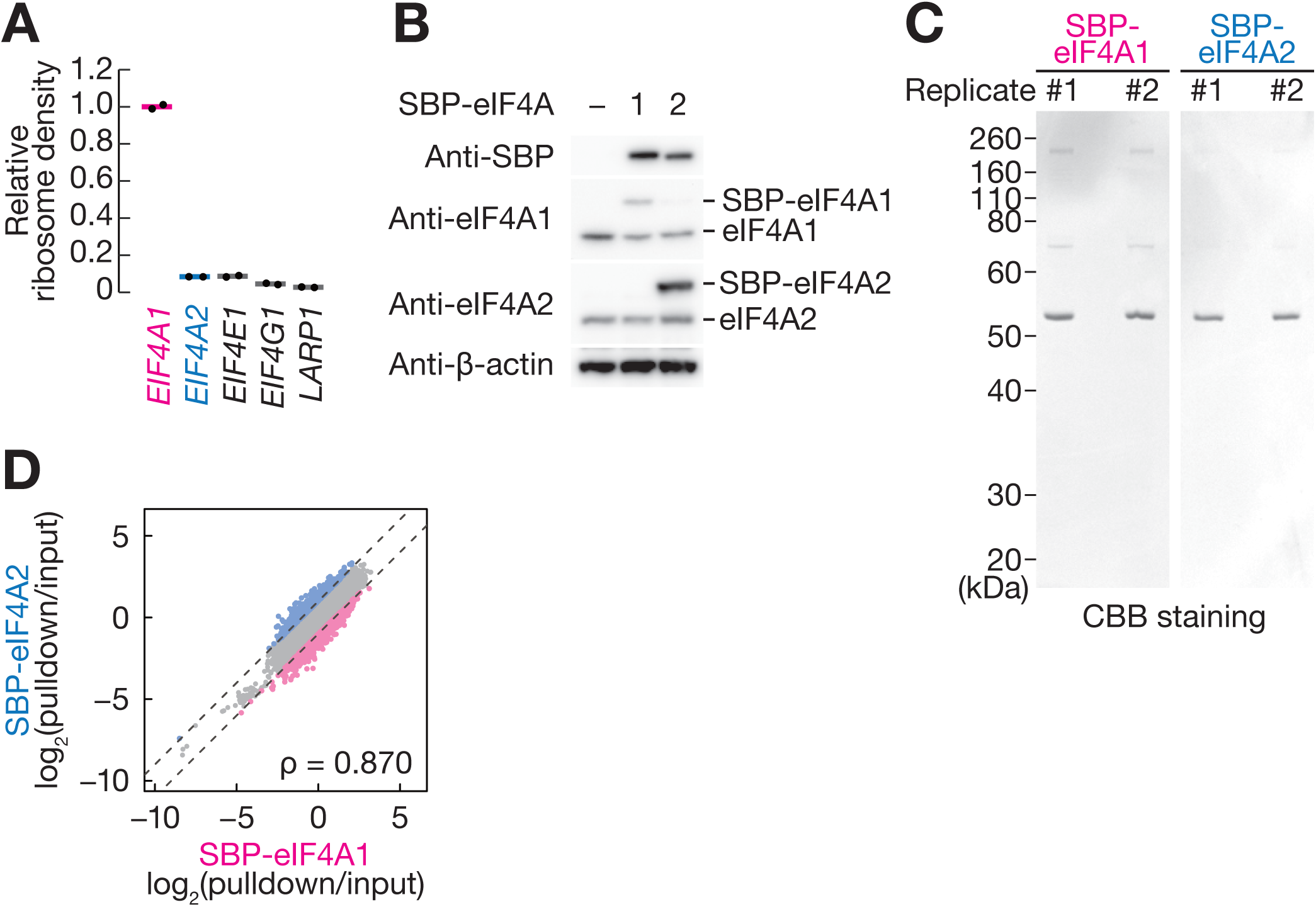
Two eIF4A paralogs show biased RNA interactions, Related to Figure 1. (A) Relative ribosome density measured by ribosome profiling of the indicated genes in HEK293T cells. (B) Western blotting of SBP-tagged eIF4A1 and eIF4A2 expressed in HEK293T cells. β- actin was used as loading control. (C) CBB staining of purified SBP-eIF4A1 and SBP-eIF4A2 proteins for RIP-Seq analysis. (D) Comparison of eIF4A1 and eIF4A2 occupancy across transcriptomes. RNAs with differential occupancies (at least twofold difference) for eIF4A1 and eIF4A2 are highlighted. ρ, Spearman’s rank correlation coefficient.

**Figure S2.**
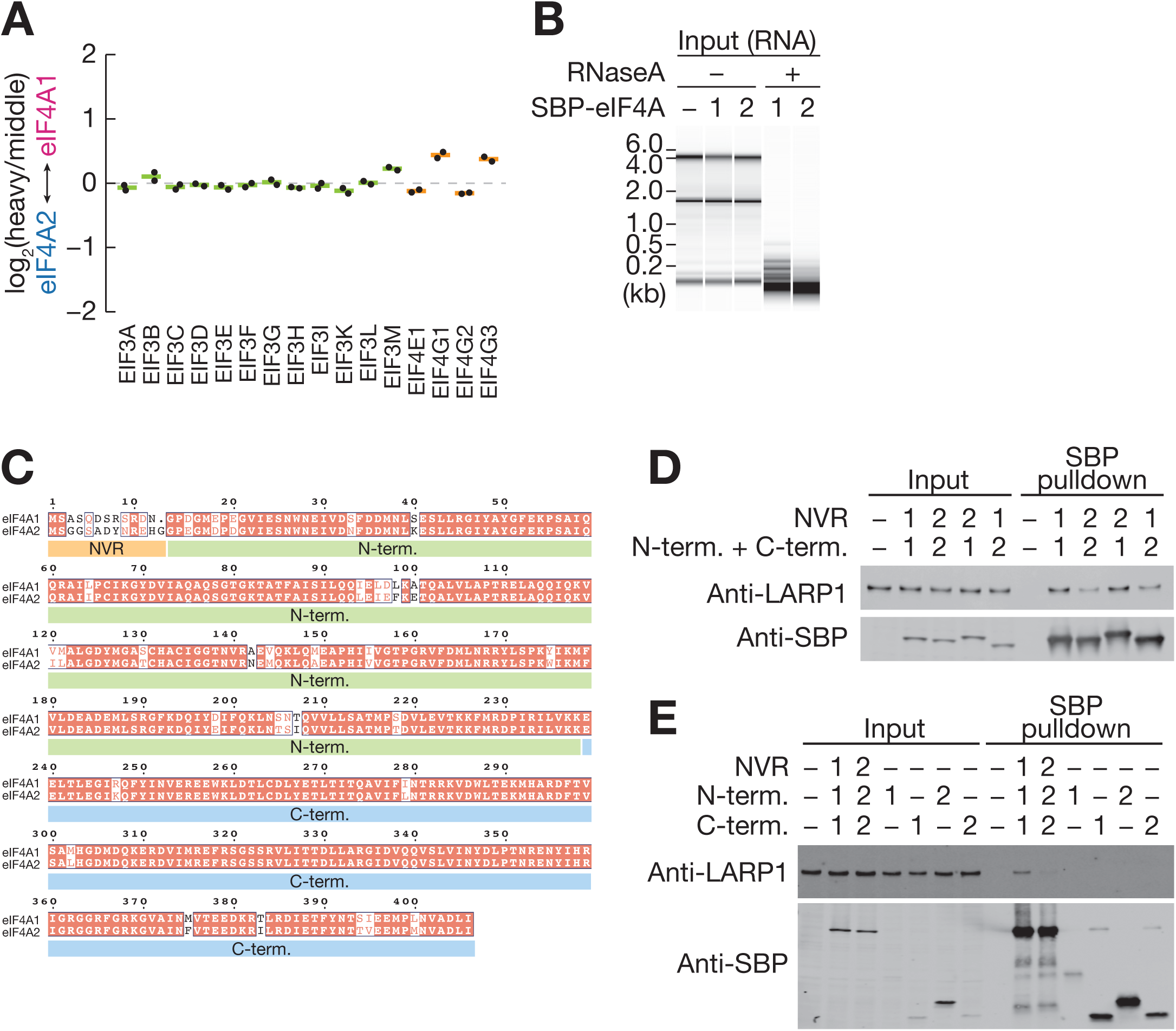
Protein binding preference for eIF4A paralogs, Related to Figure 2. (A) Biased interaction of eukaryotic initiation factors with eIF4A paralogs, measured by SILAC mass spectrometry. Data from two replicates (points) and the mean (bars) are shown. (B) Electropherogram of RNAs from lysates used for Figure 2C. (C) Sequence alignment of human eIF4A1 and eIF4A2. (D and E) Western blotting for LARP1 protein associated with engineered eIF4A proteins.

**Figure S3.**
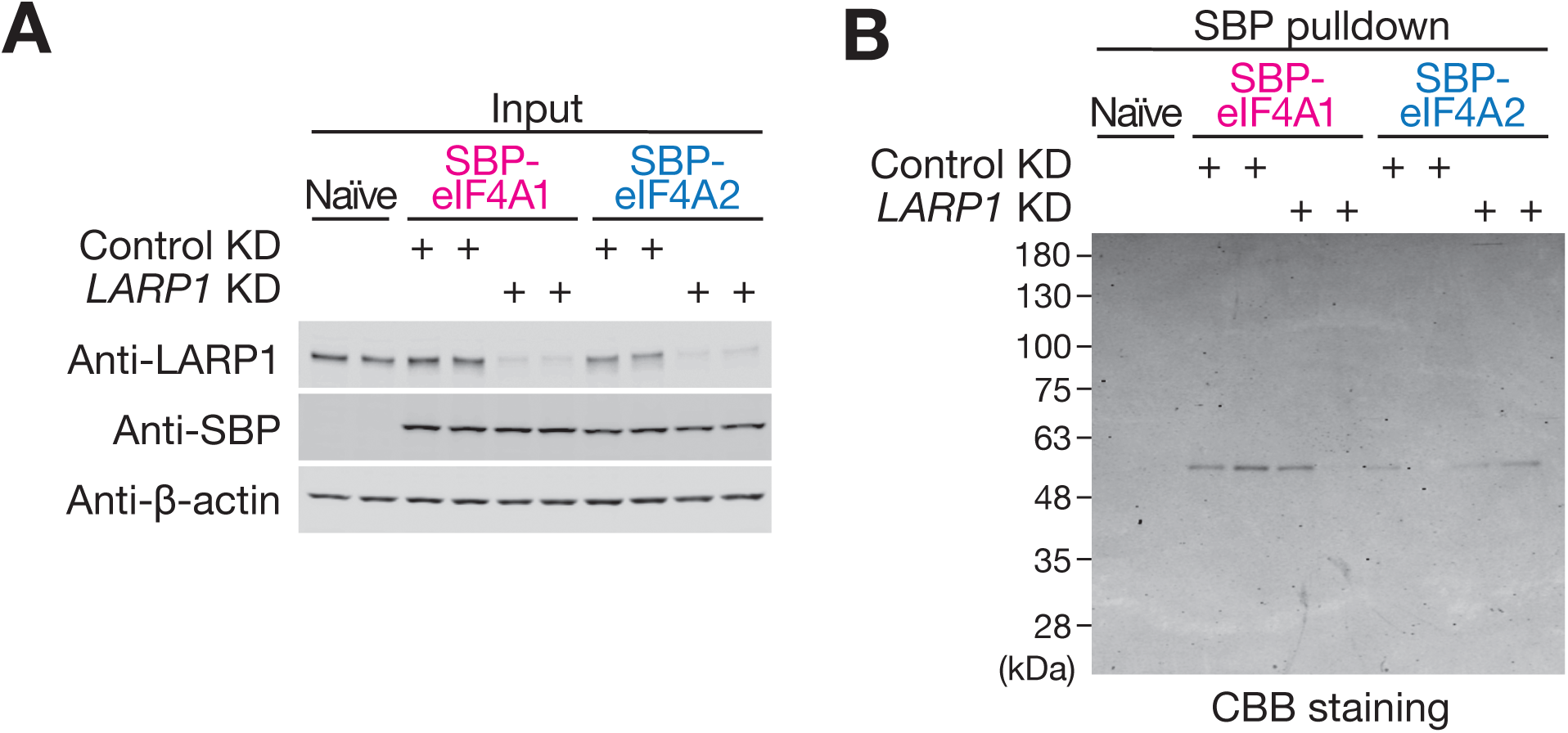
Characterization of LARP1-knockdown cells, Related to Figure 3. (A) Western blotting for LARP1 protein after knockdown. (B) CBB staining of purified SBP-eIF4A1 and SBP-eIF4A2 protein for RIP-Seq analysis from *LARP1* knockdown cells.

**Figure S4.**
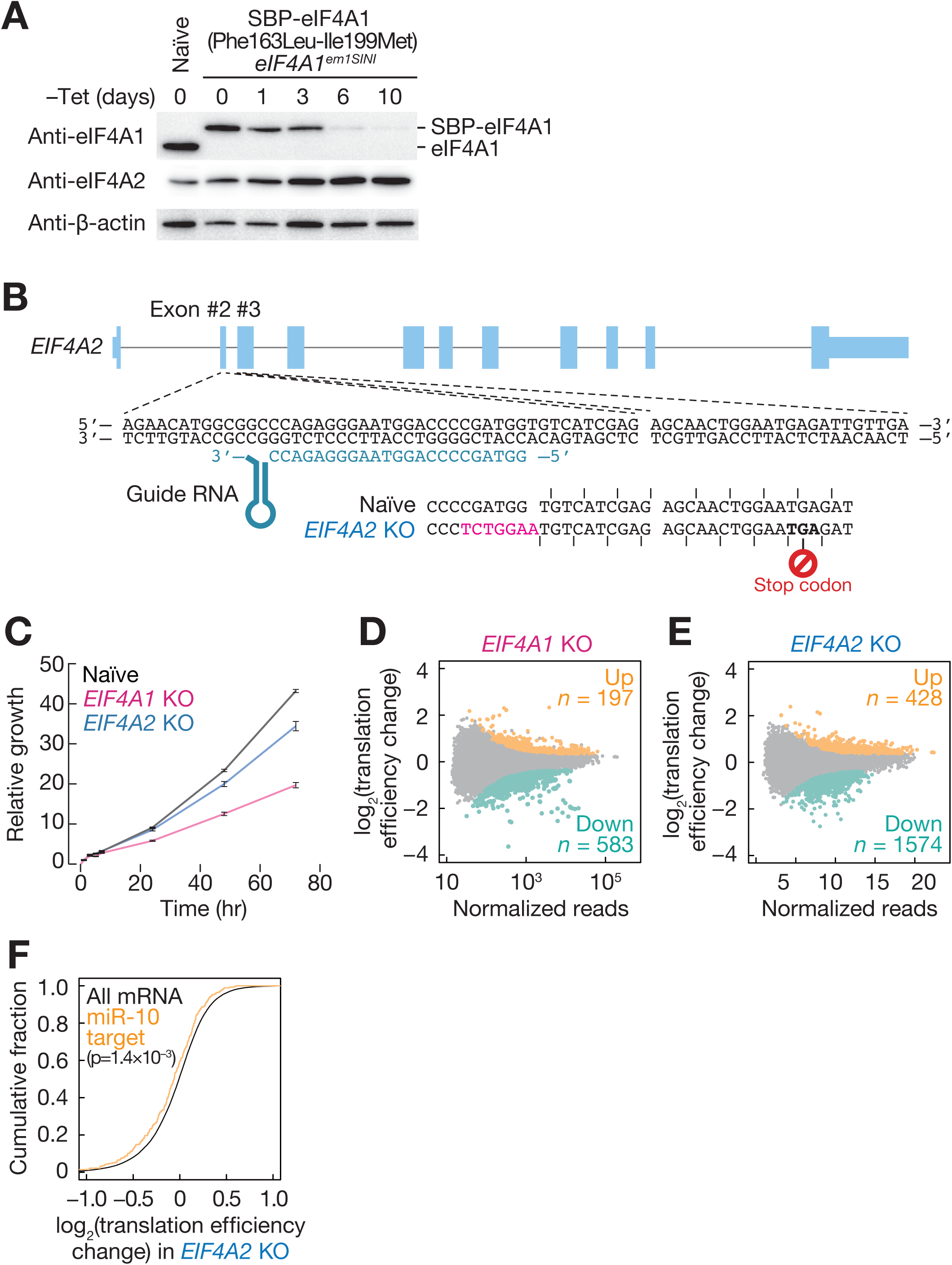
Characterization of *EIF4A1 KO* and *EIF4A2 KO* cells, Related to Figure 4. (A) Western blotting for eIF4A1 and eIF4A2 in the SBP-eIF4A1 (Phe163Leu-Ile199Met) *eIF4A1^SINI^* cell line. Cells were cultured in tetracycline-free medium. β-Actin was used as loading control. (B) Schematic of gRNA designed for *EIF4A2* gene knockout and mutation introduced by CRISPR/Cas9. (C) Growth curve of naïve, *EIF4A1* KO, and *EIF4A2* KO cells. (D and E) MA plot for the translation efficiency change in *EIF4A1* KO (D) and *EIF4A2* KO (E) cells. Upregulated and downregulated RNAs (defined as FDR < 0.05) are highlighted. (F) Cumulative distribution of all or miR-10-targeted mRNAs along the translation efficiency change in *EIF4A2* KO cells. The p-value was calculated by Mann-Whitney *U* test.

**Figure S5.**
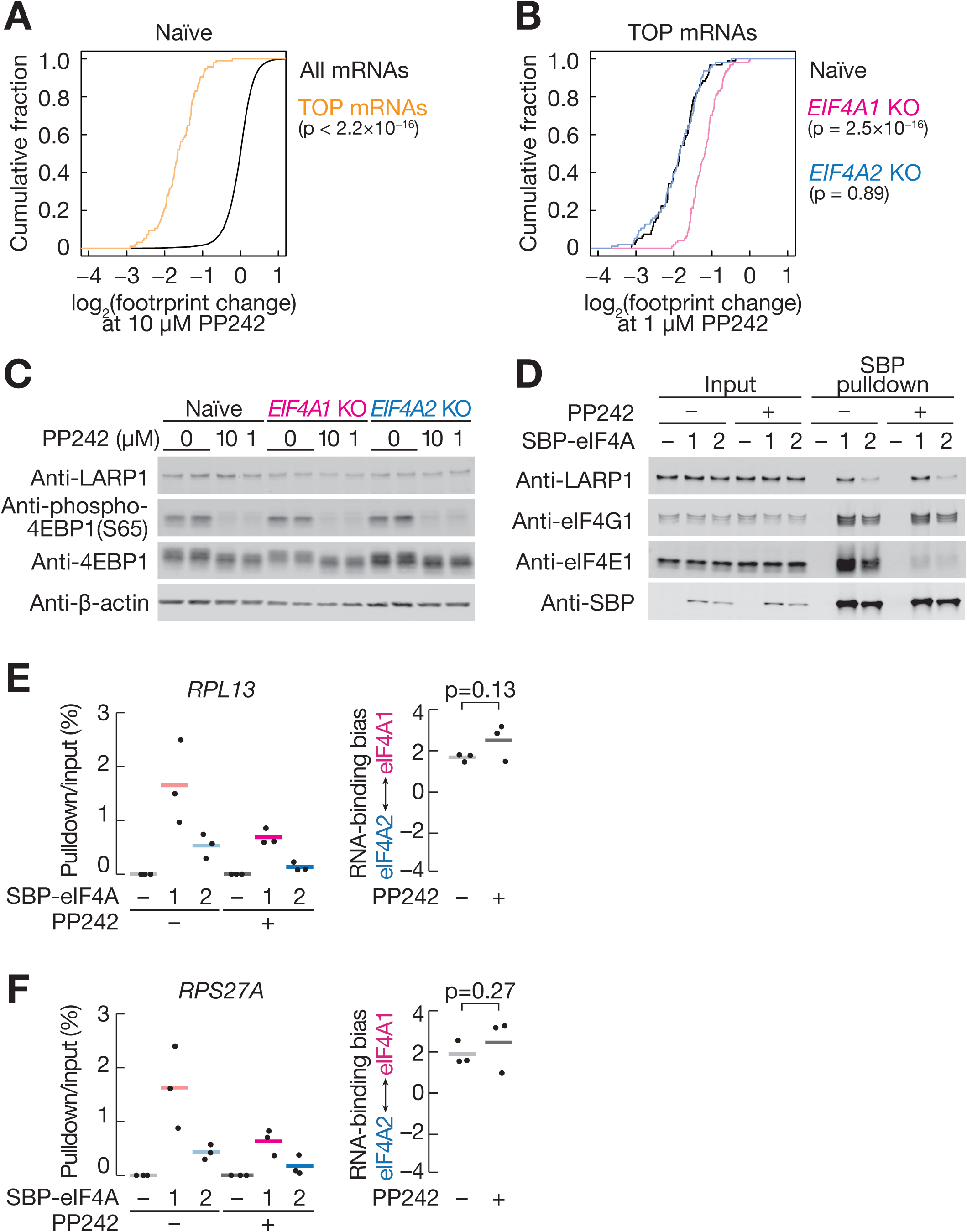
Translation repression upon mTOR inhibition is attenuated in *EIF4A1 KO* cells, Related to Figure 5. (A) Cumulative distribution of all or TOP mRNAs along translation changes upon treatment with 10 µM PP242 treatment in naïve cells. The p-value was calculated by Mann-Whitney *U* test. (B) Cumulative distribution of TOP mRNAs along translation changes upon treatment with 1 µM PP242 treatment in naïve, *EIF4A1* KO, and *EIF4A2* KO cells. The p-values were calculated by Mann-Whitney *U* test. (C) Western blotting of LARP1, 4EBP1, and phosphorylated 4EBP1 (S65) upon 1 µM and 10 µM PP242 treatment. β-actin was used as loading control. (D) Western blotting for proteins associated with SBP-tagged eIF4A proteins. Cells were treated with 10 µM PP242 before cell lysis. (E and F) Relative amounts of *RPL13* (E) and *RPS27A* (F) mRNAs associated with SBP-tagged eIF4A paralogs, measured by RT-qPCR. Cells were treated with 10 µM PP242 before cell lysis. RNA-binding bias is shown at right. Data from three replicates (points) and the mean (bars) are shown. The p-values were calculated by Student’s *t* test.

**Figure S6.**
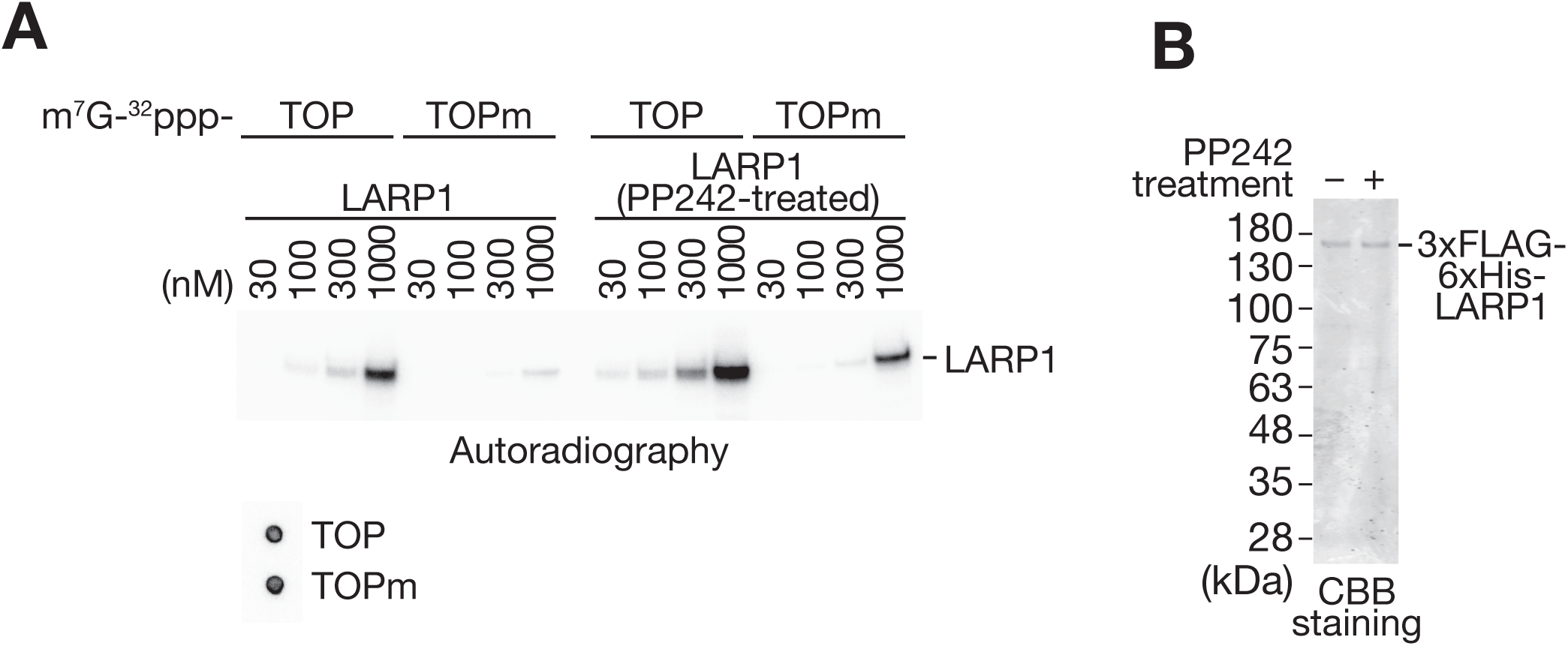
Dephosphorylation of LARP1 protein enhances the affinity to TOP motif, Related to Figure 6. (A) RNA crosslinking assay between recombinant LARP1 and 0.4 µM cap-labeled TOP or TOPm RNAs. 3×FLAG-6×His-LARP1 purified from HEK293T cells treated with or without 10 µM PP242 was used. Input cap-labeled TOP or TOPm RNAs were spotted on the nylon membrane. (B) CBB staining of recombinant 3×FLAG-6×His-LARP1 purified from HEK293T cells treated with or without 10 µM PP242.

**Table S1. List of RNAs that show biased binding to eIF4A paralogs, Related to Figure 1.**

Each transcript is listed with its Ensembl transcript ID, normalized read counts, RNA-binding bias, FDR, gene name, and gene description.

**Table S2. List of the binding ratio of each protein, Related to Figure 2.**

Each protein is listed with its UniProt protein ID, protein name and description, peptide counts; ratios of high and middle, light and high, light and middle; and FDRs for each comparison.

**Table S3. List of translation efficiency changes in *EIF4A1* KO and *EIF4A2* KO cells, Related to Figure 4.**

Each transcript is listed with its Ensembl transcript ID, normalized read counts, log2(translation efficiency-fold change to mean), FDR, gene name, and gene description for *EIF4A1* KO and *EIF4A2* KO cells.

**Table S4. List of translation changes in naïve, *EIF4A1* KO, and *EIF4A2* KO cells treated with PP242, Related to Figure 5.**

Each transcript is listed with its Ensembl transcript ID, normalized read counts, log2(footprint-fold change to mean), FDR, gene name, and gene description for naïve, *EIF4A1* KO and *EIF4A2* KO cells.

